# Effects of the overall paradigm context on intensity deviant responses in healthy subjects

**DOI:** 10.1101/2024.01.02.573901

**Authors:** Ekaterina A Yukhnovich, Kai Alter, William Sedley

## Abstract

Three experiments have been carried out to explore Mismatch Negativity responses to intensity deviants in a roving intensity deviant paradigm in control and tinnitus groups. The first experiment used interspersed blocks of two tinnitus-like frequencies set by each participant with tinnitus, which were usually around 1/3 of an octave apart. On the other hand, two later studies used interspersed blocks tones at tinnitus-like frequencies and at 1 kHz. This was the only difference in the paradigms used, however, there were differences in the patterns displayed by the control group in the first study compared to the other two. Three groups of healthy controls were recruited to measure responses to intensity deviants when different frequencies were used for the alternating blocks. For one group, the whole experiment was set at a single frequency; for the next, blocks were played at 6 kHz and at a frequency 1/3 octave below 6 kHz (small difference); the last group was presented with blocks that had tones at 6 kHz and 1 kHz frequencies (large difference). Overall, the Mismatch Negativity responses in the small difference group were opposite to the large difference and the single frequency group. It would be useful to see whether these results generalise to other experiment designs such as attended and ignored stimulus conditions, different stimulus durations, non-isochronous, or paradigms with frequency deviants.

## Introduction

Three experiments have been carried out to explore the differences between control and tinnitus group responses to intensity deviants [1-3]. The original study involved two frequencies, which were centre frequency of tinnitus and just outside of tinnitus [1]. These two frequencies were 0.37 of an octave apart, so fairly close together. The other two studies involved a tinnitus frequency (average = 5.075 kHz) and a 1 kHz frequency, which were usually quite far apart from each other (on average, 2.3 octaves) [2]. Additionally, most participants had some level of hearing loss at the tinnitus frequency but not at the 1 kHz frequency. There were some differences in the responses seen in the paradigm with only tinnitus-like tones compared to the paradigm that included 1 kHz, despite no other differences being present.

The crucial differences between the two 1 kHz-inclusive studies were seen in the MMN response patterns to intensity deviants at the tinnitus frequency stimuli in the control group. In the original study, responses to the edge tinnitus frequency were larger in magnitude to the centre tinnitus frequency but otherwise showed the same pattern, so the edge frequency was used in the subsequent studies, and therefore the focus of the comparison for this introduction will be on high (i.e. tinnitus-related) frequency responses (Table 1). The MMN pattern was also similar between the two frequencies in the two later studies, though possibly somewhat underpowered in the second study [2] compared to third [3]. The paradigms in the three studies were identical, except for the addition of the 1 kHz tone instead of a second tinnitus-like tone. However, this seemed to affect the pattern of responses in the control group, with the amplitudes inverting with the addition of the more distant other frequency (i.e. 1 kHz). Additionally, unpublished data, where intensity deviant responses in healthy participants were elicited in paradigms with frequency deviants only or paradigms including both frequency and intensity deviants, showed different patterns of response to downward deviant (DDs) and upward deviants (UDs) for intensity.

**Table 1.**
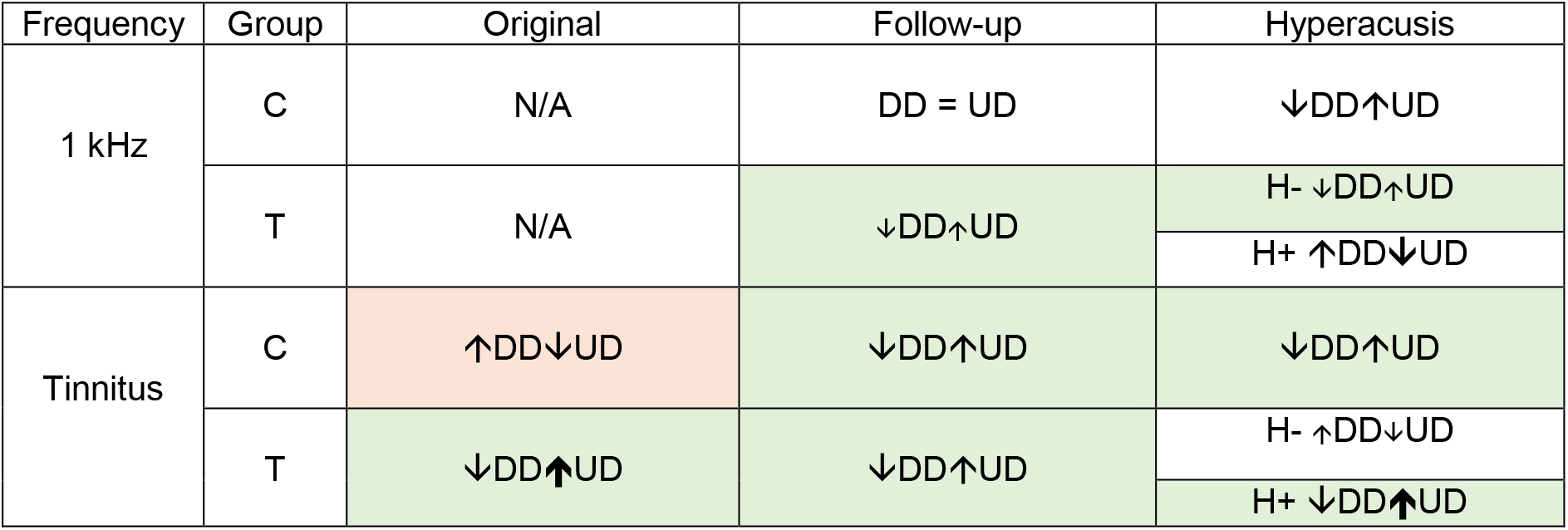
Pattern of responses in each group to downward deviants and upward deviants. C = controls, T = tinnitus group. The hyperacusis study also includes H- (without hyperacusis) and H+ (with hyperacusis).↑ = stronger response, ↓ = weaker response. The arrows are smaller when the directionality was not as pronounced and thicker when the differences were driven by a particular deviant. Green colour shows the major similarities in responses between the three studies, while red indicates the main difference in findings between the paradigms.

| Frequency | Group | Original | Follow-up | Hyperacusis |
| --- | --- | --- | --- | --- |
| 1 kHz | C | N/A | DD = UD | ↓DD↑UD |
|  | T | N/A | ↓DD↑UD | H- ↓DD↑UD<br>H+ ↑DD↓UD |
| Tinnitus | C | ↑DD↓UD | ↓DD↑UD | ↓DD↑UD |
|  | T | ↓DD↑UD | ↓DD↑UD | H- ↑DD↓UD<br>H+ ↓DD↑UD |

The impression emerging from the differences between these similar studies, mostly in the control group, is that the inclusion of frequency changes elsewhere in the experiment may affect intensity mismatch asymmetry within a single frequency. Potential mechanisms responsible could be adaptive processes in which neural responses adapt to stimuli in a particular environment through mechanisms such as frequency-specific adaptation or contrast gain control (a mechanism in which neuronal responsiveness to intensity is dynamically adjusted based on the global context/statistics of recent stimulation) [4-7]. For example, bat auditory neurons react differently to a target sound in the context of being preceded by certain sound sequences [8]. Similar processes have been studied in the visual and somatosensory systems, where they help, for example, to adjust to a new context of a changing light intensity through enhancement or suppression [9, 10]. Gain control may be an important part of ensuring efficiency of the auditory neurons, e.g. they may have a redundancy-reducing effect within a local sensory environment [4, 11]. However, the mechanisms of auditory gain control more generally or its effects on perception are not fully understood [5].

A previous experiment was conducted to see whether different variability levels between auditory stimuli would affect neuronal responsiveness [4]. To achieve this, the experimenters played tone sequences with three contrasts (low: -/+5 dB deviant (relatively constant), medium: -/+ 10 dB and high: -/+15 dB (high variability)). They found that responses to a standard sound depended on the overall context of the paradigm in two ways. Firstly, gain non-linearly increased as stimulus contrast decreased, thus partially compensating for the smaller changes through a more sensitive neural firing rate. Contrast gain is a form of statistical adaptation in the auditory cortex that creates representations of the complex statistical environment and the change in this environment over time [12]. It involves dynamic range adaptation of neurons within the auditory system and, based on the representation of contrast in the environment that has been created over time, predicts the way in which neurons will need to adjust their gain [12]. Secondly, decreases in neural gain occurred faster (in ms) than increases in neural gain. Therefore, adaptation to a high-contrast context was faster than to a low-contrast context. Additionally, neural gain was best modulated within their responsive frequency range, though further frequencies also had an effect on gain, and if mean level of the stimuli was low, overall effectiveness of this mechanism was reduced. Extracellular recording of A1 neurons experiment has similarly studied contextual interactions in a two-tone paradigm [10] and showed that if a 6 kHz tone was presented after a 7 kHz tone, response to the 6 kHz tone was suppressed, but it was not suppressed if presented after a 2 kHz tone, as the first tone was not in the receptive field of the second tone. Interestingly, when the researchers compared the suppression of response to a louder second tone (80 dB) versus a quieter second tone (30 dB), the region of suppression was larger for the quieter second tone than the loud one. Additionally, the firing rate evoked by the louder second sounds was higher than for quieter second tones. Therefore, suppression was proportional to intensity of the first tone and inversely proportional to intensity of the second tone, as well as being related to the frequency difference between the first and second stimuli.

It has been previously concluded, in support of the aforementioned studies, that through the process of forward masking, frequency tuning of auditory cortical neurons could be dynamically modulated by preceding stimuli, particularly by neurons with similar characteristic frequencies and particularly at higher stimulus levels [9]. These studies taken together may indicate that both frequency and intensity of the tones played in roving paradigms may interact to affect response to intensity deviants, and inter-frequency effects may be stronger in paradigms with small frequency contrast contexts than large frequency contrast contexts.

Therefore, the present experiment was carried out in order to establish how difference in intensity mismatch asymmetry (IMA) to the roving intensity paradigm result from frequency differences present between blocks within the whole experimental paradigm. Specifically, to compare either a single frequency, two similar frequencies, or two far-apart frequencies. Based on the previous results, hypotheses were:

- The paradigm with a small frequency difference will create an opposite IMA pattern to the paradigm with a large frequency difference: larger downward and smaller upward deviants for small frequency differences, and smaller downward and larger upwards deviants for large frequency differences;
- Based on the unpublished data from our group, the single-frequency paradigm may show a prolonged P50 and an attenuated N100 unlike the paradigms that include two frequencies.

## Materials & Methods

### Participants

Participants were randomly assigned to one of three groups, who were presented a roving oddball intensity paradigm featuring either: 1) all stimuli at a single frequency (N=15), 2) blocks alternating between two frequencies, with a small frequency difference (1/3 of an octave) (N=15), or 3) blocks alternating between two frequencies, with a large frequency difference (1 kHz and 6 kHz) (N=13). Participants only did one version of the experiment because 1) all three paradigms in one session would have blurred any specific context effects, and 2) on separate days, there might have still been some carryover or familiarity effects from previous session/s. Participants were recruited from affiliated volunteer lists at Newcastle University. Inclusion criteria included being over 18 years old, and being able to make an informed choice about volunteering. Exclusion criteria included presence of any type of tinnitus (subjective, objective or intermittent), using ongoing sedating or nerve-acting medications, and mental health conditions severe enough to interfere with everyday life activities. It was also ensured that there were no significant group differences in age or pure-tone audiometry results. Approval was given by the Newcastle University Research Ethics Committee, and all participants gave written informed consent according to the Declaration of Helsinki (reference number 5619/2020).

### Common methods: psychophysics and EEG

The psychophysical assessment in which sound intensity played during the EEG recording were determined, and the experimental design, closely followed the procedure in [2] and [1]. The main exception was the lack of tinnitus frequency due to the nature of participants. Instead, all participants listened to 6 kHz tones. In the small frequency difference group, participants also listened to tones 1/3 of an octave lower than 6 kHz (4.643 kHz) as, on average, the edge tinnitus frequency in the original study was 1/3 of an octave lower than tinnitus centre frequency. In the large frequency difference group, the other tone was 1 kHz, to replicate the paradigm context of the two later IMA studies.

### Statistical analysis

Statistical analysis was performed using MATLAB. To compare evoked responses between the three groups at the ‘tinnitus’ frequency (6kHz), a two-way ANOVA was used, with subject group and intensity as the main factors of interest, along with the interaction terms. Post-hoc analysis included Tukey Honest Significant Tests to determine any significant differences between the ERP amplitudes of the three groups.

## Results

Table 2 shows means and standard errors (SE) of the demographic information of the 3 groups. One-way ANOVAs were carried out to see whether age or HQ scores were different between the three groups. Age and HQ scores were not significantly different between the groups (p=0.843 and p=0.790, respectively). Average HQ scores were also below the cut-off score for hyperacusis presence. Chi-square test showed that there were significant differences in gender between groups (p=0.01). As the pure tone audiometry results were not normally distributed at the majority of frequencies/ears Kruskall-Wallis tests were performed and showed that across all three groups, there were no significant differences in the hearing ability at 0.5, 1, 2, 4, 6 or 8 kHz in right or left ears. At 0.25 kHz, there was also no difference between the groups in the right ear, but there was a significant difference in the left ear (p=0.035). This was acceptable as this frequency was not used as a stimulus. Importantly, at 6 kHz there were no significant differences between the three groups: p=0.560 for the left ear and p=0.300 for the right ear.

**Table 2.** Descriptive statistics of the three study groups. Means are standard errors are given to every group for their age and HQ scores, as well as the gender split.

|  | Single Frequency | Small Difference | Large Difference |
| --- | --- | --- | --- |
| Age | 48.40 (4.90) | 48.87 (4.51) | 52.15 (5.03) |
| Gender | 11 f, 4 m | 12 f, 3 m | 7 f, 6 m |
| HQ Scores | 9.87 (1.66) | 8.87 (1.33) | 10.38 (1.75) |

### Time course of the stimulus response

Grand average ERP data for channel FCz (with P9/P10 reference) across standard and deviant responses for all stimulus conditions and in each group was used to attempt to determine timeframes for quantifying P50, N100, and MMN responses, based on visual inspection (Fig 1). To calculate the MMN difference waveform, standard responses were subtracted from their equivalent deviant conditions.

**Fig 1.**
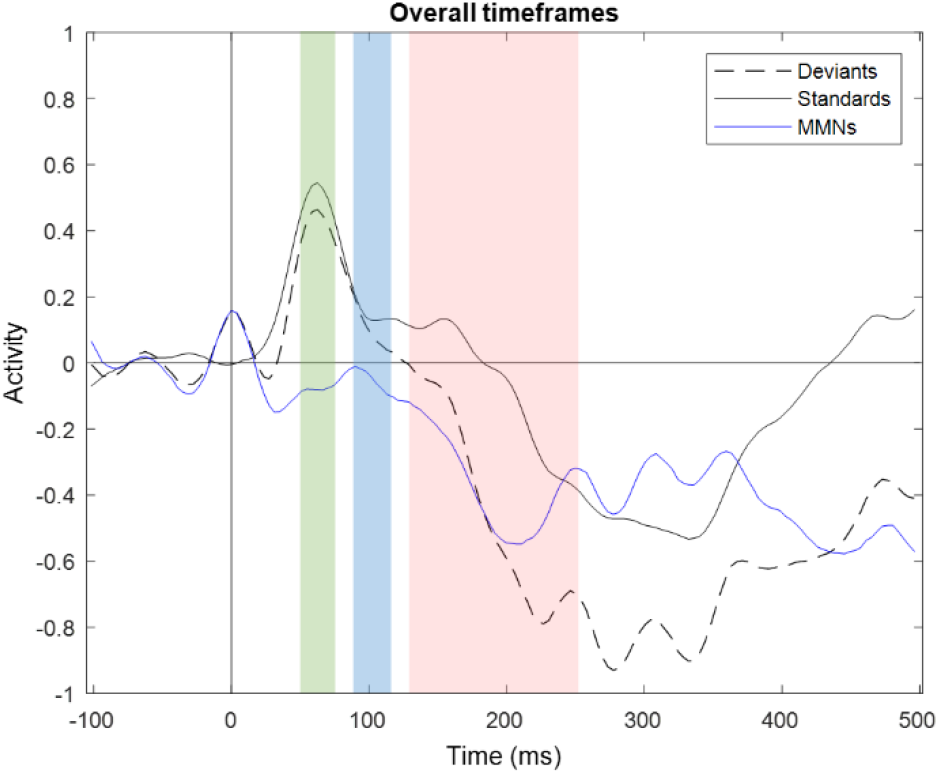
Standard, deviant and MMN waveforms broadly combined across all groups at 6 kHz. The chosen P50 timeframe was 50-75 ms (green), N100 was 90-110 ms (blue). The MMN timeframe was 140-250 ms (red).

### Standard and pure deviant waveforms

The average waveforms of responses to 6 kHz tones of each group are shown on Fig 2.

**Fig 2.**
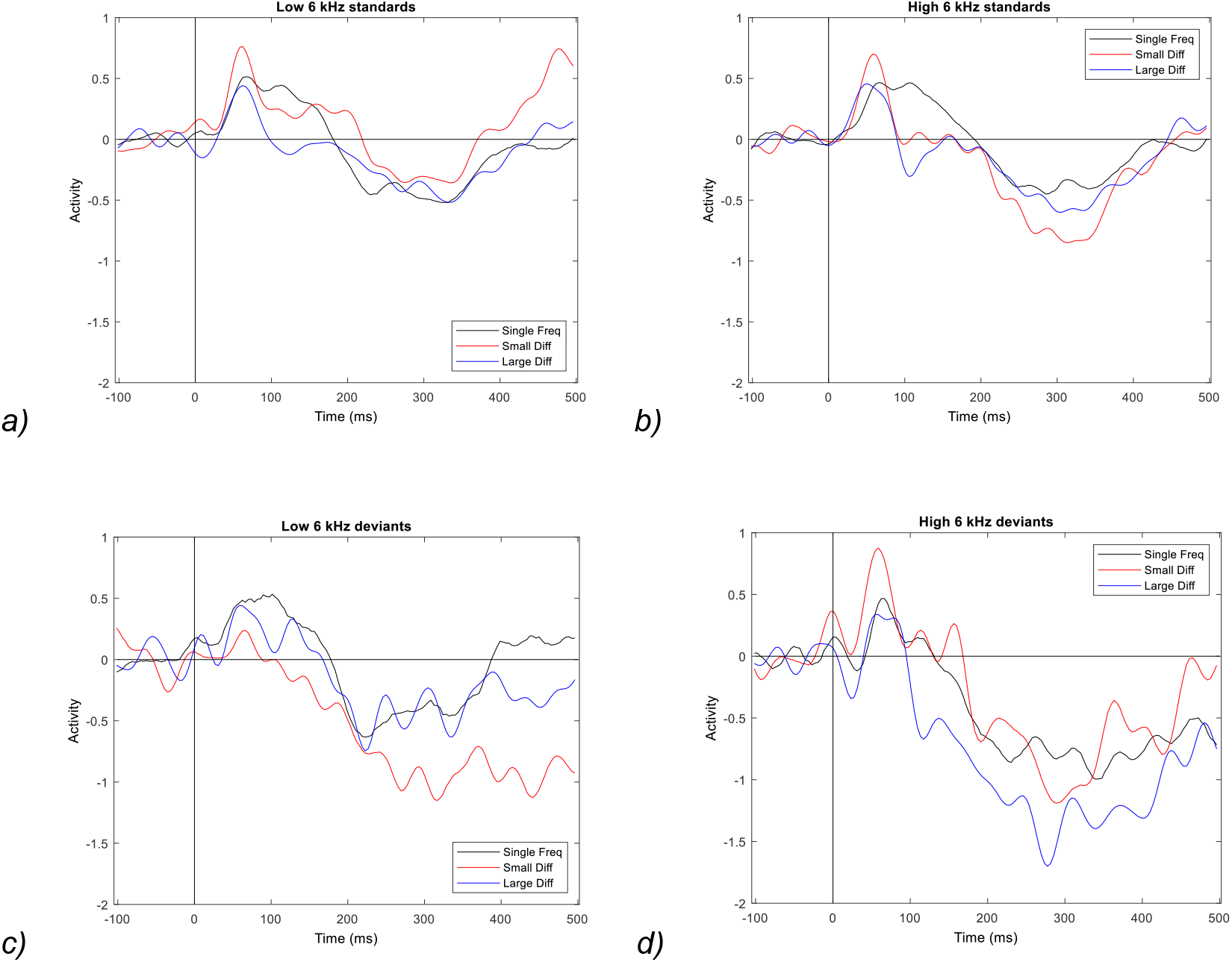
Average waveforms to standard and deviant conditions at 6 kHz. On the left, are the quieter stimuli conditions and on the right are the louder stimuli conditions. Responses to standard tones are shown on a) and b), while responses to raw deviant stimuli are shown on c) and d). Single frequency group is shown in black, small difference group is shown in red, and large difference group is shown in blue.

### P50 responses

Fig 3 shows P50 responses to the two standard and two raw stimuli. A two-way ANOVA (group, intensity) showed a main group effect on P50 amplitudes in response to standard stimuli. The small difference group had significantly larger responses than both of the other groups at both intensities (all p<0.001).

**Fig 3.**
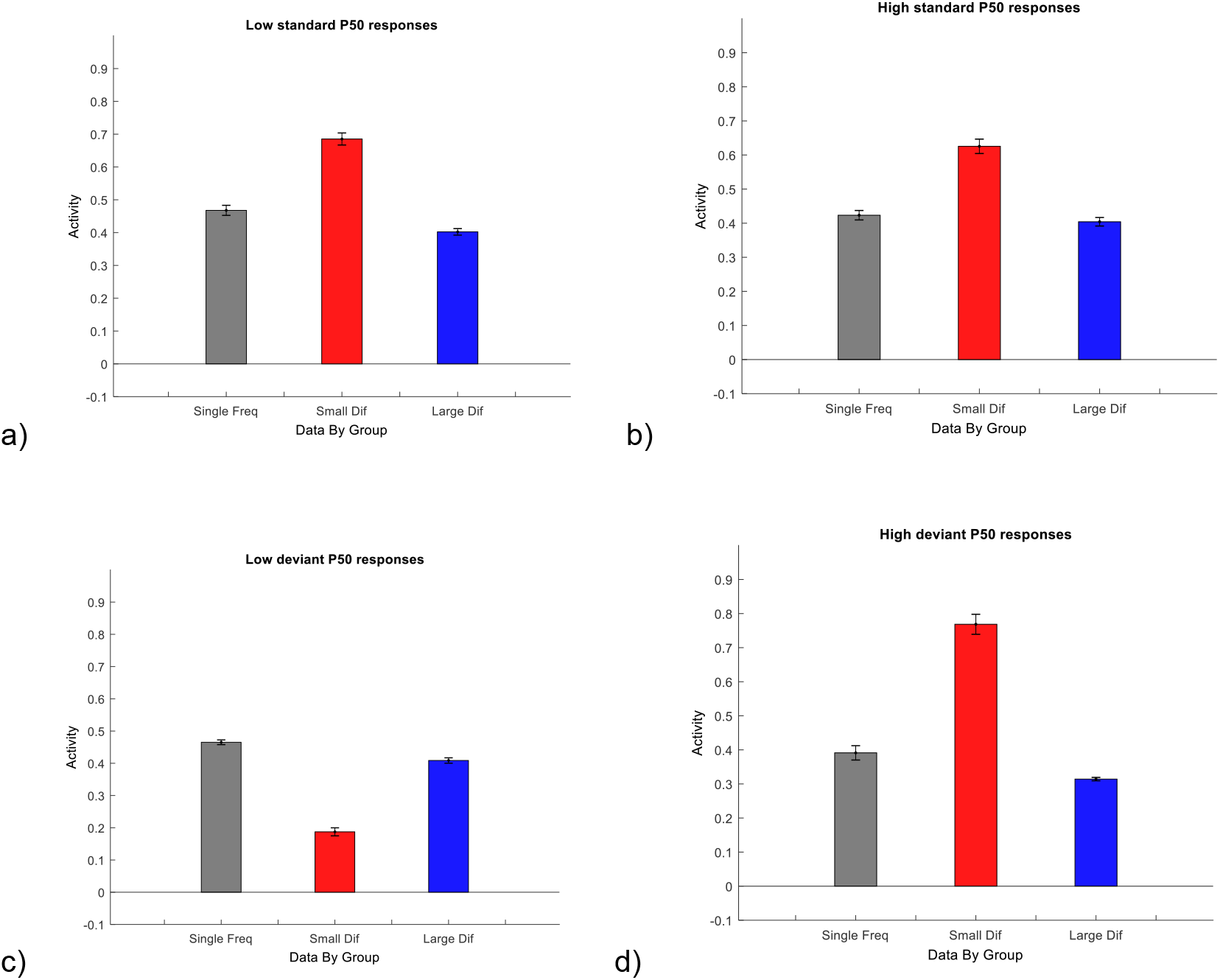
Standard and deviant responses in the P50 timeframe. Top two charts represent averaged responses to standard stimuli bottom charts represent averaged responses to raw deviant stimuli. The order of the groups in all charts is: single frequency (grey), small frequency difference (red), and large frequency difference (blue).

A two-way ANOVA (group, intensity) showed main effects of both factors on P50 response amplitudes to deviant tones, as well as a significant interaction effect between group and intensity. Post-hoc Tukey test showed that the single frequency group had larger amplitudes compared particularly to small difference group (p<0.001), but also large difference group (p=0.025). In response to high deviant stimuli, the small difference group had significantly larger amplitudes than both other groups (p<0.001), but no differences were seen between single frequency and large frequency groups. This was a striking finding, as while the single frequency and large difference groups had relatively similar responses to both deviants, small difference group had a very clear pattern. The small difference group had weaker P50 amplitude to low deviants than the other two groups, and stronger P50 responses to high deviants than the other two groups.

### N100 responses

Fig 4 shows N100 responses to two standard and two raw stimuli. A two-way ANOVA (group, intensity) showed main effects of both main factors on N100 amplitude in response to standard tones (p<0.001). This is not surprising, as the single frequency group did not seem to display an N100 (Fig 1). The N100 amplitudes were significantly weaker in response to low standard tones than high standard tones both in the small difference group (p<0.001) and large difference group (p=0.006), though the large difference group showed stronger N100 responses to standard stimuli overall.

**Fig 4.**
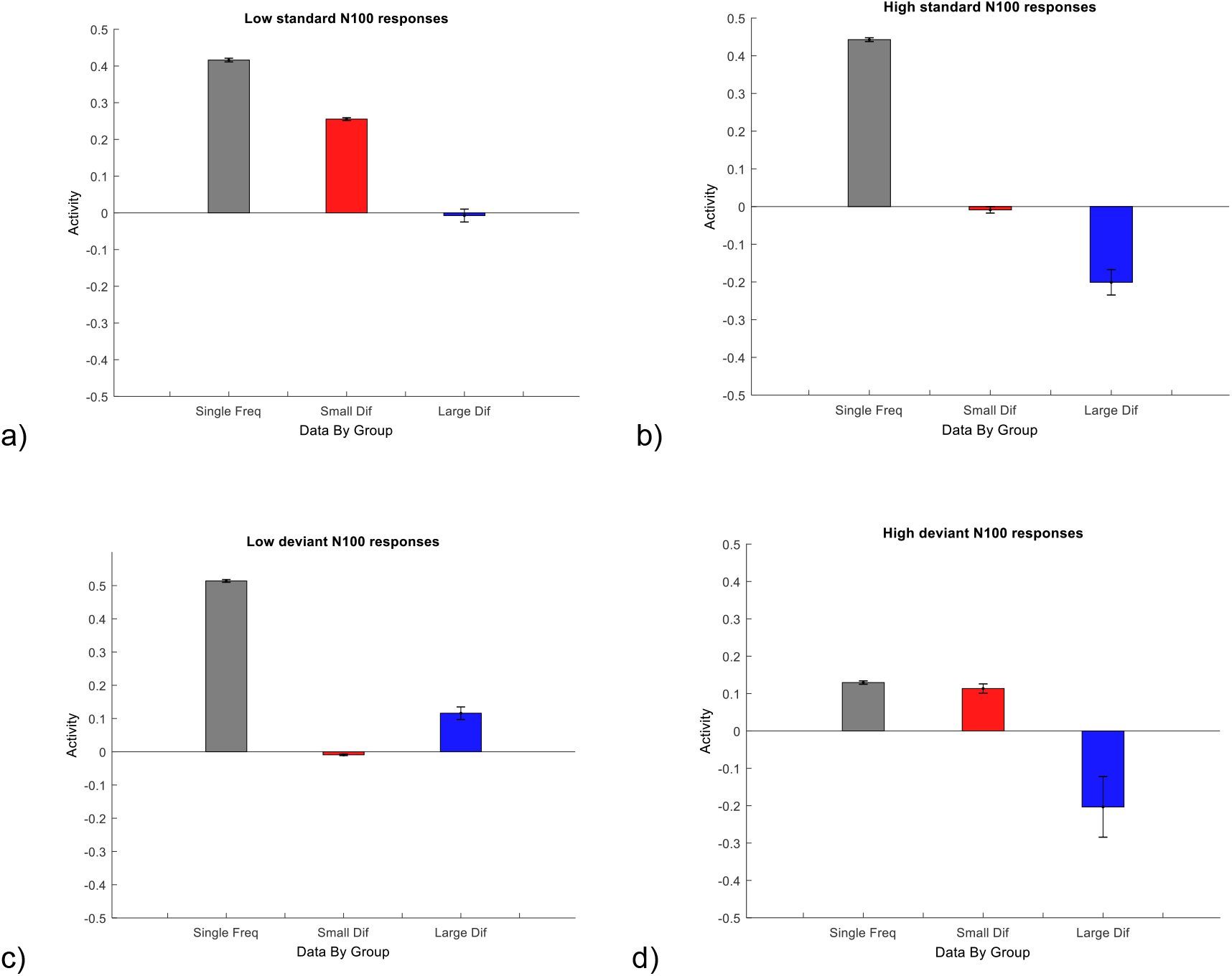
Standard and deviant responses in the N100 timeframe. Top two charts represent averaged responses to standard stimuli bottom charts represent averaged responses to raw deviant stimuli. The order of the groups in all charts is: single frequency (grey), small frequency difference (red), and large frequency difference (blue).

A two-way ANOVA (group, intensity) also showed main effects of both main factors on N100 amplitude in response to deviant tones, as well as a significant interact between the factors (all p<0.001). Tukey post hoc tests showed that in response low deviant tones, the single frequency group had significantly more positive amplitudes during the N100 timeframe as expected (p<0.001), but the small difference group had significantly lower amplitude than the large difference group (p<0.001). In response to high deviant tones, the large difference group had a significantly stronger N100 compared to the other two groups (p=0.011 compared to single frequency and p=0.015 compared to small difference).

### MMN responses

The average waveforms of responses to 6 kHz tones of each group are shown on Fig 5. Fig 6 shows MMN responses to the two deviant conditions. A two-way ANOVA (group, intensity) showed that both main factors have significant effects on the amplitude of MMN responses, with a significant interaction between the factors (all p<0.001). Post hoc Tukey tests showed that in response to DDs, single frequency and large difference groups produce responses that are not significantly different from each other (p=0.890), however both of these are significantly weaker than the response produced by the small difference group (both p<0.001). In response to UDs, the large difference group produced an MMN response that was significantly stronger than both single frequency and small difference groups (p<0.001). However, the single frequency group still produced a significantly stronger response than the small difference group (p<0.001).

**Fig 6.**
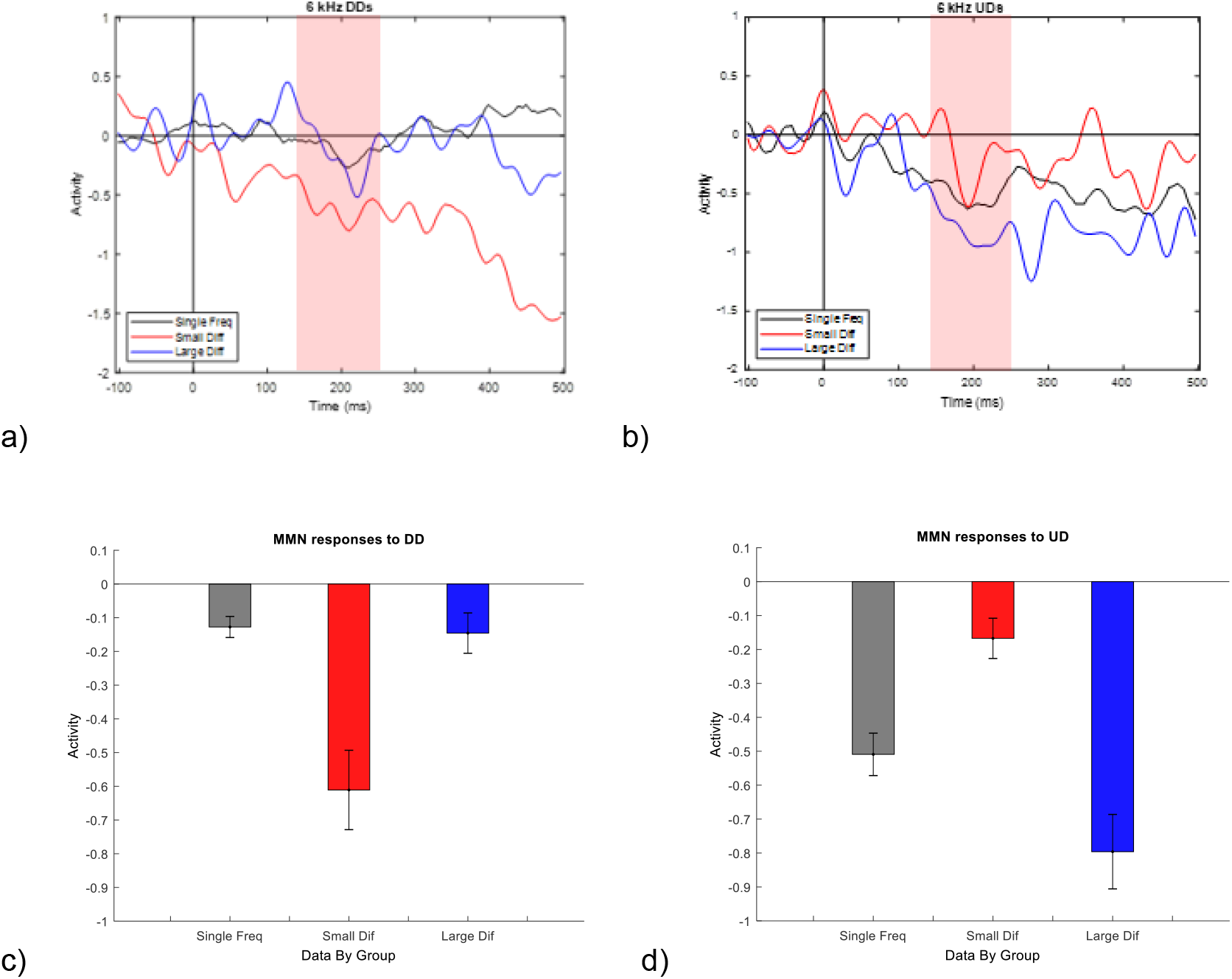
MMN responses. Top two charts represent Average MMN waveforms for each subject group, in each condition at 6 kHz. The MMN timeframe was 140-250 ms (red). On the left, is the downward deviant and on the right is the upward deviant. Single frequency group is shown in black, small difference group is shown in red, and large difference group is shown in blue. Bottom two charts represent averaged responses to downward deviants and upward deviants. The order of the groups in all charts is: single frequency (grey), small frequency difference (red), and large frequency difference (blue).

## Discussion

### Summary

Overall, the deviant/MMN responses in the small difference group were often opposite to the large difference group. The large difference group somewhat followed the waveform patterns seen in the single frequency group. The small frequency difference group MMN was stronger in response to DDs than UDs, while the other two groups show the opposite pattern of stronger MMN to upward intensity deviants.

P50 amplitudes were the most evident in the small difference paradigm, with an interesting pattern coming through in response to raw deviant stimuli. The low deviant responses in the small difference paradigm were lower than the other two groups, while the high deviant responses were larger than the other two groups. Single frequency and large difference paradigm responses remained similar within the two intensity conditions. N100 responses were the clearest in the large difference paradigm (possibly due to the lower P50 amplitude compared to small difference). In response to raw low deviant stimuli, N100 seemed attenuated in all paradigms. All three groups showed a negativity around 200-400 ms post standard stimuli. In the single frequency paradigm, in lieu of the completely attenuated N100 responses, P50s were prolonged.

### Possible underlying mechanisms

Previous research points towards habituation likely playing a large part in the formation of the waveforms in the current findings. In repetition suppression experiments, N100 amplitude has been previously found to rapidly reduce even in response to the second stimulus in the train and continue to attenuate in the following four stimuli; P50 amplitude also decreased after the first stimulus in this paradigm, but not between the second stimulus and later stimuli [13]. Further, the N100 data was entirely consistent with other unpublished data within the lab of the current authors, showing that long single-frequency paradigms completely attenuate the N100 response. It is instead replaced with a positive response that extends the P50. It may be explained, based on published literature, as the effect of extreme repetition positivity.

Interestingly, P50 response to standards was least attenuated in the small frequency difference group but the N100 was least attenuated in the large difference group. It could be expected that the small difference paradigm responses may be more sensitive to changes in stimulation due to increased contrast gain [4]. This may be true for P50 responses, except the low deviants, as on average the pre-attentive reaction to both standard and deviant stimuli was stronger than in the other two paradigms. In the low deviants, the suggestion that suppression is inversely proportional to the intensity of a second tone especially within responsive frequency fields may somewhat relate to the reduced P50 and N100 in the small difference group [10]. A detailed study on different combinations of stimuli patterns and their ability to create reproducible fundamental differences in topologically constrained small network responses concluded that networks create stable responses after repeated stimulation through various plasticity effects [14]. If such differences can be created in small networks, it could be postulated that such differences could carry into larger scale networks. However, studies performed on a small scale could show the effects of two similar frequencies in the paradigm in unexpected ways compared to whole brain studies. For example, different results were found for intracranial vs EEG in terms of N100 latencies [13]. Nonetheless, in a previous EEG study, researchers presented 115 trains of unique frequency stimuli, with one condition being wide range of frequency, and another being narrow range of frequency. They found that N100 amplitude was dependent on the frequencies presented during an experiment, with neurons adjusting to the range of stimulation history dynamically, potentially through lateral inhibition [6]. This seems similar to the dynamic range adaptation in intensity paradigms. Overall, it is possible that the differences between the paradigms seen in the current study are fairly fundamental to the auditory system. The single frequency and large difference paradigms may show may similar responses because the field affected by changes in gain does not reach the lower frequency. This can explain the discrepant findings between the original IMA study [1] and the following related studies [2, 3].

Lower N100 suppression has been related to over-inclusion of background sounds, meaning that participants with more N100 (in the current study, large difference group) were more aware of background sounds, which the previous researchers attributed to passive attention switching towards novel sounds [15]. Within their study, the authors found that a stronger P50 suppression was related to a stronger MMN response for intensity deviants (though to 1 kHz stimuli). This seems true in the current study as well, for example, the raw deviant P50 amplitude was inversely related to MMN amplitudes, e.g. small difference group had weaker P50 but stronger MMN amplitudes in response to DDs, and the other way round for UDs. The association between P50 and MMN amplitudes was also present in and earlier EEG study [3], where T+H+ had a particularly decreased P50 amplitude in response to a high deviant and had a particularly strong MMN for UD, while T+H-had a similar pattern in response to low deviant and DD. An interesting hypothesis was suggested, in which the P50 suppression was related to filtering out of irrelevant stimuli and thus allowing the brain to more efficiently detect changes in the stimulus, leading to a larger MMN [15]. Accordingly, in a round-about way, it is possible that the small difference paradigm is still creating more sensitive responses even to DDs.

While habituation has been a guiding theory, some authors argue that the inter-stimulus interval length effect on N100 is more in line with the refractoriness hypothesis than habituation [16]. Refractoriness hypothesis suggests than N100 decrease may be due to recovery period of particular neural generators, and not due to learning of the repeating stimuli. The two accounts differ in their predictions for the waveform in response to the stimulus after the deviant tone: the refractoriness hypothesis assumes no response recovery after a deviant tone, no exponential attenuation of N100 and no N100 amplitude change after and ISI longer than the recovery period [17]. On the other hand, the habitation hypothesis suggests continuous N100 decrease and dishabituation once a deviant tone or even tonal language, such as spoken Mandarin, are played [18]. In the current data, after particularly repetitive stimuli (single frequency group), the N100 was completely gone. Additionally, the authors state that refractoriness refers only to the immediate past prior to the current stimulus. Therefore, the different results based on the overall paradigm context would not quite fit this hypothesis. Notably, the current data cannot disprove either hypothesis as while there was no N100 in the standard in response to the single frequency paradigm, N100 was not present in response to the deviants either, and the ISI used in this study was much shorter than the recovery period [19]. However, it may be interesting to pay closer attention to the behaviour of the waveform throughout the length of the experiment and particularly at and just after deviant stimuli.

### Future directions

Importantly, the observations discussed in here apply to the particular set of experimental parameters tested. There are, however, other paradigm specifics that could affect the shape of the overall waveform (particularly N100 and P200 ERPs), such as length of inter-stimulus intervals (ISI) and attention [20, 21].For example, temporal attention can alter cortical gain, and gain in neurons responsible for predictive error signals [22]. This could potentially be due to optimisation of predictions about the sensory input, as one MEG study showed a significant difference between expected vs unexpected tones in attended condition, but not unattended condition, when the expectation was based on a prior cue that specified the instructions for every experimental block [21]. It would be useful to see how they generalise to other experiment designs such as attended and ignored stimulus conditions, different stimulus durations, ISIs, non-isochronous, paradigms with frequency deviants, etc.

## Conclusions

Overall, the wider context of the different frequency ranges in the experimental paradigms dramatically influenced both P50 and intensity mismatch responses in a reciprocal way. Unchanging or widely separated frequencies resulted in upward intensity deviant MMNs being larger than for downward, and narrowly separated frequencies produced the opposite pattern. This effect applies even when the frequency changes are remote (e.g. tens of seconds to minutes away) from the intensity responses they influence. It is likely that similar influences occur (logically, and from group’s unpublished data) when frequency changes are recent or ongoing from the intensity responses they influence.

Based on results of the previous research within the lab of the current authors [3], hyperacusis (with or without tinnitus) was characterised by increased upward and reduced downward intensity MMNs, the pattern related to hyperacusis might be best revealed using narrowly different frequencies, to give the maximal contrast between the intensity conditions. Conversely, tinnitus without hyperacusis was characterised by increased downward intensity deviants, and therefore may be best revealed using single-frequency or widely-spaced frequency differences.

In summary, tinnitus and hyperacusis show distinct signatures in intensity deviant MMN response profiles, as do control responses depending on the range of frequencies used in the paradigm, and future studies should consider optimising their paradigm for the condition under study based on these observations.

